# Mechanistic insights into RAN translation within the Huntingtin coding sequence

**DOI:** 10.1101/2025.09.02.673827

**Authors:** Amy Krans, Richard M. Albertson, Connor J. Maltby, Peter K. Todd

## Abstract

Nucleotide repeat expansions support production of toxic peptides from putatively non-coding regions of the human genome without use of an AUG initiation codon. This process, known as RAN translation, was originally described at CAG repeats with products generated in all three possible reading frames. Recently, cryptic mis-splicing of mRNAs in which the CAG repeat acts as a splice acceptor was proposed as an alternative mechanism for RAN protein production. To investigate this, we generated DNA and RNA based reporters to assess RAN translation in the context of a CAG repeat expansion in exon 1 of the Huntingtin gene (*HTT),* associated with Huntington Disease. HTT CAG repeats support RAN translation in both the alanine (GCA) and glutamine (CAG) frames, with or without upstream AUG start codons or near-AUG cognate codons, and at levels comparable to RAN translation from CGG and GGGGCC repeats. CAG RAN translation products were readily detectable from *in vitro* transcribed RNAs transfected into human cells and rodent neurons - suggesting that plasmid based aberrant splicing into CAG repeats cannot fully explain the observed phenomena. These findings indicate that RAN translation from HTT CAG repeats shares key mechanistic parameters with other disease-associated repeat expansions.

**Key Points:**

- Translation at HTT CAG repeats occurs in at least two reading frames absent any AUG start codon.
- Like RAN translation at other repeats, HTT CAG RAN translation is enhanced by cell stress.
- HTT CAG RAN translation readily occurs from in vitro transcribed RNAs in neurons.

## INTRODUCTION

Huntington disease (HD) is an age-related neurodegenerative disorder characterized by psychiatric and neurological symptoms including psychosis and dementia in combination with a progressive movement disorder featuring chorea and gait difficulties(1). The disease is inexorably progressive and almost uniformly lethal. HD is caused by a CAG repeat expansion in exon 1 of the Huntingtin gene, *HTT* (2). CAG repeat expansions of 36 or more are considered pathogenic, with larger repeats resulting in an earlier age of onset and more rapid progression(3,4). The CAG repeat track is translated into a polyglutamine stretch within the mutant HTT protein (mHTT), which makes the protein more prone to aggregation and triggers the formation of neuronal intranuclear inclusions and alters proteostasis(5). The CAG repeat expansion also impacts the normal functions of the HTT protein, leading to epigenetic and transcriptomic alterations which manifest as changes in synaptic function and cellular fitness(6).

Numerous nucleotide repeat expansions that cause neurodegenerative disease support a novel form of translational initiation known as repeat-associated non-AUG (RAN) translation(7–9). This process was first described at CAG repeats in the context of the opposite strand RNA generated in Spinocerebellar Ataxia type 8 – where removal of an AUG codon just 5’ to the CAG repeat did not preclude translation of protein from the repeat(9). Subsequently, RAN translation has been observed in genes with CGG, CCG, CUG, CCUG, GGGGCC, CCCCGG, and CAG repeats and across a number of coding and non-coding sequence contexts(9–17). RAN translation products can be generated in multiple reading frames and from antisense transcripts in genes where bidirectional transcription occurs(18,19). Some of these protein products are toxic when expressed absent the repeat sequence(20,21) and are found in aggregates in patient brains(10,11,18).

An interesting observation about RAN translation is that it can occur in any part of a gene. RAN translation in Fragile X-associated Tremor/Ataxia Syndrome (FXTAS) occurs in the 5’ untranslated region of *FMR1,* where the repeat is thought to stall scanning ribosomal pre-initiation complexes and allow for initiation at near-AUG cognate codons (such as ACG or GUG) to produce a polyglycine-containing protein, FMRpolyG(13,22). At the same time, translation also occurs less efficiently in the poly-alanine frame – which lacks any near-AUG codon and appears to initiate within the repeat itself (23). For both reading frames, initiation is largely dependent on the 5’ m^7^G cap and is blocked by inhibition of the RNA helicase eIF4A – both of which suggest a scanning mode of initiation(22). In contrast, *C9orf72* associated Amyotrophic Lateral Sclerosis and Frontotemporal Dementia (C9 ALS/FTD) results from a GGGGCC hexanucleotide repeat expansion in the first intron of the gene, where it is less obvious as to how the repeat might engage with ribosomes(24,25). Some data supports a model where scanning ribosomes engage with GGGGCC repeats through repeat “exonization” (cryptic transcription initiation upstream of the repeat and splicing from cryptic sites downstream of the repeat that together place the repeat element in the 5’ leader of C9orf72) (26–29), placing it in a similar sequence context to the CGG repeat in *FMR1*. Consistent with this finding is evidence supporting a role for a CUG codon just 5’ to the repeat in the glycine- alanine (GA) reading frame that appears important for its translation (26,29) and potentially for translation in other reading frames. Alternatively, the repeat may serve as an internal ribosomal entry site (IRES) that allows for 5’ m^7^G cap-independent translation to generate dipeptide repeat (DPR) containing proteins – potentially from lariat RNAs that escape degradation and transit to the cytoplasm (25).

In 2015, RAN translation was identified as a potential pathogenic mechanism in Huntington Disease. Bañez-Coronel and colleagues observed production of polyalanine and polyserine products from plasmid-based reporters lacking any AUG start codon(30). Furthermore, they detected potential RAN translated proteins in patient brains using antibodies raised against their predicted carboxyl termini – which would be out of frame with the main polyglutamine reading frame of HTT. RAN proteins were more abundant in juvenile cases with larger repeats. Consistent with this, BAC transgenic mice carrying a CAG repeat (but not an interrupted CAGCAA repeat) exhibited RAN translation products late in their disease course(31). However, subsequent work failed to observe generation of such products in HD KI mice(32).

Recent findings also suggest that RAN translation products generated from plasmid-based reporters with CAG repeats might be alternatively explained by cryptic splice events within the plasmid that would create mRNAs with an AUG codon just 5’ to the repeat that would support AUG-initiated translation(33). Another consideration is that the CAG codon is embedded within a protein coding region, it would be expected to engage predominantly with actively translating ribosomes rather than pre-initiation complexes – different from both CGG repeats in *FMR1* and GGGGCC repeats in *C9orf72*. Indeed, some evidence suggests that out-of-frame products might be generated instead by ribosomal frameshifts from the glutamine reading frame into serine and alanine reading frames(34–36).

As a comparative mechanistic assessment of how RAN translation might occur in the context of the HTT locus has not been conducted, we generated a series of reporter agents for RAN translation from HTT CAG repeats in different sequence contexts and compared this process to RAN translation reporters with CGG or GGGGCC repeats. We observe RAN translation in the polyalanine and polyglutamine, but not consistently the polyserine, reading frames of the HTT CAG repeat- consistent with some but not all prior work on CAG RAN translation(15,30,32,37). Production of the glutamine RAN translation products does not require an AUG or near-AUG cognate codon while the alanine RAN translation products relies largely on a CUG near-cognate codon. Initiation remains largely 5’ m^7^G cap-dependent and it is significantly enhanced by integrated stress response activation in some reading frames, shared features to RAN translation in CGG and GGGGCC repeats. We did not observe any evidence supporting plasmid rearrangement or splicing as an explanation for the observed products(33). Further supporting this, RAN translation from CAG repeats was also detected from RNAs generated via *in vitro* transcription in *in vitro* cell lysates, and transduced HEK293 cells and rodent neurons. Together, these data provide insights into the unusual process of RAN translation and helps to harmonize data discrepancies in the field.

## METHODS

### Plasmids

The AUG-NL-3x FLAG, GGG-NL-3x FLAG, FMR1 polyG and polyA, and C9orf72 GA and GP plasmids have been described previously. (19,22) A cassette with 3 stop codons in each reading frame was inserted upstream of the multiple cloning site of pcDNA3.1(+) using SacI-HF and NheI-HF restriction enzymes. None of these plasmids contain AUG codons in any repeat reading frame 3’ to the CMV promoter or T7 transcription start site.

Initial WT HTT and RAN HTT plasmids were a gift from Dr. William Yang (UCLA) and contained 140 and 123 CAG repeats, respectively. WT HTT plasmid contained all of exon 1 (including the 5’UTR) of the HTT gene with the conical AUG codons at positions 1 and 8 in the protein. The RAN HTT plasmid began at the coding sequence of HTT with the conical AUG codons at positions 1 and 8 mutated to UGA and AUC, respectively, and continued through exon 1. Because the WT and RAN plasmids did not have the same 5’ sequence, the 5’UTR sequence that was present in the WT HTT plasmid was inserted into the RAN plasmid using EcoRI and StuI restriction enzymes.

A triple tag cassette containing a 3x FLAG, 3x HA, and 5x Myc tag, one tag for each reading frame, was generated by GeneWiz (Figure 1A). The cassette was inserted downstream of the WT HTT or RAN HTT plasmid using EcoRV and PspOMI restriction sites. The resulting triple tag plasmids (WT TT and RAN TT) had 3x HA-tagged glutamine (polyQ), 5x Myc-tagged serine (polyS), and 3x FLAG-tagged alanine (polyA). For controls, the HA tag and Myc tags were fused to the C-terminal end of GFP using complementary annealing oligonucleotides. For the addition of both tags, the restriction enzymes BsrGI and PspOMI were used. The Myc tag was inserted using the following oligo set, 5’- GTACAAGGAACAAAAACTCATCTCAGAAGAGGATCTGTAAG-3’ and 5’- GGCCCTTACAGATCCTCTTCTGAGATGAGTTTTTGTTCCTT-3’. The HA tag was inserted using the following oligos: 5’-GTACAAGTACCCATACGATGTTCCAGATTACGCTTAAG-3’ and 5’- GGCCCTTAAGCGTAATCTGGAACATCGTATGGGTACTT-3’. The oligos were resuspended in 100mM KOAc, 30mM HEPES such that the OD_260_/100μL was roughly equal for each pair. The oligos were mixed at equal volumes and heated at 95°C for 5min and then allowed to cool down for 1hr. While the oligos were cooling, the GFP containing plasmid was digested with BsrGI and PspOMI at 37°C. After 1hr, the digested plasmid and annealed oligos were ligated together using the Rapid DNA Ligation kit (Sigma) following the manufacturer’s protocol.

**Figure 1:**
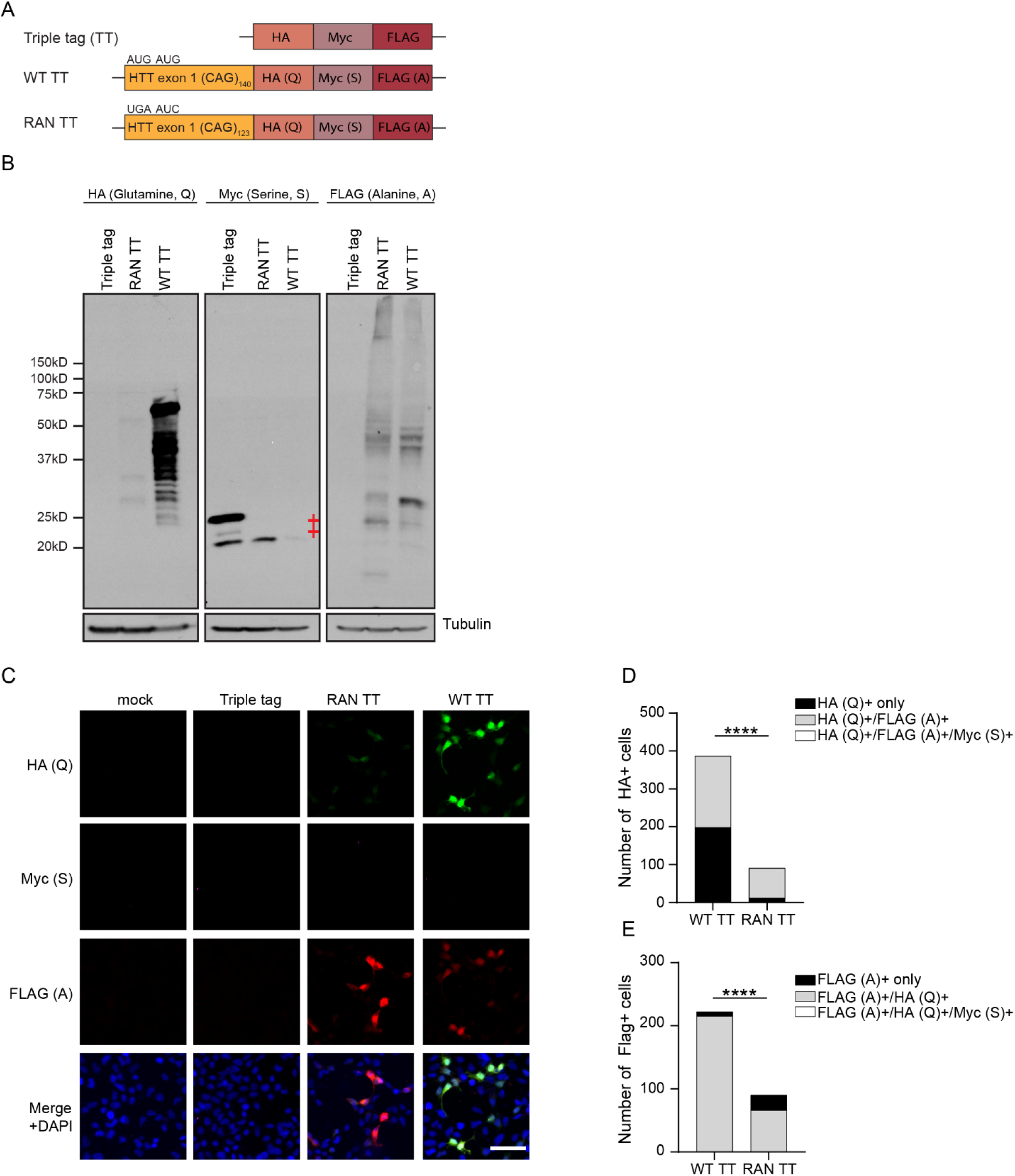
HTT reporters demonstrate alanine and glutamine frame RAN translation products. A. Schematic of HTT reporters with triple tag cassette. B. Representative western blot images of triple tag reporters from HEK293 cells. Tubulin was used as a loading control. ‡ denotes non-specific band associated with Myc antibody. C. Representative ICC images from HEK293 cells expressing triple tag reporters; HA (glutamine) is green, Myc (serine) is magenta, and FLAG (alanine) is red. DAPI was used as a nuclear counterstain. Scale bar = 50μm. D. Coincident expression of alanine (FLAG+) and serine (Myc+) reading frame products in glutamine (HA+) positive cells transduced with WT or RAN triple tag reporters. E. Coincident expression of glutamine (HA+) and serine (Myc+) reading frame products in alanine (FLAG+) positive cells transduced with WT or RAN-only reporters. For D and E, data expressed as total number of cells across two independent experiments. **** p<0.0001 two-sided Chi square test comparison of proportions.

A fluorescent GFP tag was inserted in place of the triple tag cassette using the same restriction sites as above (Sup Fig 1F). GFP was PCR amplified using 3 different forward primers (Q: 5’- TACTACTACGATATCGGCCGGGGTGAGCAAGGGCGA-3’, S: 5’- TACTACTACGATATCGCCGGGGTGAGCAAGGGCGA-3’, A: 5’- TACTACTACGATATCCCGGGGTGAGCAAGGGCGA-3’) to generate plasmids in each reading frame. In addition to inserting the frame shift with the forward primer, the AUG of GFP was mutated to a GGG so the fluorescent signal observed would be from the upstream HTT sequence. The same reverse primer was used for all sets (5’-GTAGTAGTAGGGCCCTTTACTTGTACAGCTCGTCCATG).

WT and RAN HTT were inserted into GGG-NL-3x FLAG pcDNA3.1 using HindIII/EcoRV and EcoRI/EcoRV restriction site pairs, respectively, (RAN or WT Q). A cassette of 67 interrupted CAGCAA repeats (coding 134 polyglutamine, synthesized by GeneWiz) was inserted upstream of GGG-NL-3x FLAG (Scr Q_134_). Q5 site directed mutagenesis was used to insert 1 or 2 nucleotides to shift the reading frame to either serine (RAN or WT S) or alanine (RAN or WT A), respectively. Sequences for mutational analysis were synthesized by GeneWiz and inserted using StuI and EcoRI restriction enzymes. Firefly luciferase (FF) was PCR amplified from pGL4.13 and inserted into pcDNA3.1 using HindIII and XbaI restriction enzymes.

### Cell culture and transfections

HEK293 and HEK293T cells were purchased from American Type Culture Collection. Cells were cultured in Dulbecco’s modified Eagle’s medium (DMEM) with 10% fetal bovine serum and 1% penicillin-streptomycin at 37°C with 5% CO_2_.

All mammalian rodent work was approved by the Institutional Animal Care and Use Committee (IACUC) at the University of Michigan. Rat primary hippocampal neurons were dissected from P0-P1 Sprague-Dawley rat pups. Hippocampi were isolated and dissociated using papain as previously described in Sutton et al(38). Cells were plated in media consisting of Neurobasal-A, 1x GlutaMax, and 1x B27 Supplement and maintained at 37°C, 5% CO2.

For NL assays, HEK293 cells were plated at 2x10^4^ cells/well in a 96 well plate and transfected the next day. Rat primary hippocampal neurons were plated at 3.5x10^4^ cells/well in a 96 well plate on day of dissection and transfected 7-8 days later. For DNA plasmid transfections, HEK293 cells were transfected with 50ng of NL plasmid and 50ng Firefly luciferase (FF) using FuGENE HD (Promega) according to the manufacturer’s protocol. *In vitro* transcribed RNAs (100fmoles each NL and FF) were transfected into HEK293 cells or primary neurons using Lipofectamine MessengerMax according to the manufacturer’s protocol. Luciferase assays were performed 24hrs after transfection as previously described. (19,22)

For activation of the integrated stress response, HEK293 cells were plated as described above. Cells were transfected with Viafect (Promega) for 1hr followed by treatment with vehicle (DMSO or PBS), 2μM (final) thapsigargin (TG) in DMSO, or 20μM (final) sodium arsenite (SA) in PBS for 5hrs. Cells were then harvested for NL assay.

For western blot analysis and RNA isolation, HEK293(T) cells (2x10^5^ cells/well) or rat primary hippocampal neurons (3.5x10^5^ cells/well) were plated on a 12 well plate. Cells were transfected with 500ng of HTT (NL or triple tag) plasmid DNA using FuGENE HD or 500ng *in vitro* transcribed RNA using TransIT mRNA transfection kit (Mirus), both according to the manufacturers’ protocols. Cells were harvested for analysis the day following transfection.

For immunocytochemistry, HEK293 cells (1x10^5^ cells/well) were plated on a 4-well chamber slides. The next day, cells were transfected with 500ng NL, GFP, or triple tag plasmid DNA. After 24hrs, cells were fixed and stained as described below.

### NanoLuc luciferase assay

Cells were lysed in 70μL Glo Lysis Buffer (Promega) for 5min at room temperature on a rocker. HEK293 cell lysates (25μL) were transferred to a 96 well black plate (Corning, 3915). Primary rodent neuron lysates (25μL) were transferred to a 96 well white plate (Corning, 3912). Either 25μL of NanoGlo substrate diluted in NanoGlo Buffer (for NL signal, Promega) or 25μL of OneGlo (for FF signal, Promega) solution was added to the plate. When cells were treated with thapsigargin or sodium arsenite, cell viability was also measured using CellTiter-Glo (Promega) solution in a 1:1 ratio between solution and lysate. After 5min incubation at room temperature, the plates were read on a GloMax Luminometer.

### Immunoblotting

HEK293 cell lysates were collected as follows. Media was removed and cells were washed once in 1x PBS. Cells were lysed in 200μL RIPA buffer with protease inhibitors and sonicated. Lysates were reduced in 6x SDS sample loading dye with beta-mercaptoethanol and boiled at 95°C for 5min. Samples were stored at -80°C if not used immediately.

Proteins were separated on 12% SDS-PAGE gels run at 180-200V until the dye line had run off the bottom of the gel. The gels were then transferred to PVDF membrane at 320mA for 90min at 4°C. Membranes were blocked with 5% non-fat dried milk in Tris Buffered Saline plus Tween-20 (TBST).

Antibodies were diluted in 5% non-fat dried milk in TBST and incubated overnight at 4°C, generally. Membranes were washed 3x, 10min each, with TBST and then incubated with horseradish peroxidase-conjugated or IRDye secondary antibodies for 1hr at room temperature. Proteins were either visualized on film or on an Odyssey Imaging System.

For subcellular fractionation, HEK293 cells were plated and transfected as described above. Cells were harvested after 24hrs and were washed twice with 1x PBS. Cells were incubated on ice in a hypotonic buffer (20mM Tris-HCl pH7.4, 10mM NaCl, and 3mM MgCl_2_). After 15min, 10% NP40 was added and the sample vortexed. The hypotonic fraction was collected after centrifugation at 3000rpm at 4°C. The remaining pellet was further lysed in cell extraction buffer (10mM Tris-HCl pH7.4, 2mM Na_3_VO_4_, 100mM NaCl, 1% Triton-X 100, 1mM EDTA, 10% glycerol, 1mM EGTA, 0.1% SDS, 1mM NaF, 0.5% deoxycholate, 20mM Na_4_P_2_O_7_) for 30mim on ice. The nuclear fraction was collected after centrifugation at 14000 x g at 4°C. The final insoluble pellet was resuspended in 8M urea.

### Immunocytochemistry

HEK293 cells were plated on 4 well chamber slides. The following day, cells were transfected with 500ng of DNA plasmid. After 24hrs or 48hrs, cells were fixed with 4% paraformaldehyde in PBS, permeabilizing with 0.1% Triton, and blocked with 5% normal goat serum all in PBS-MC (PBS with 1mM MgCl_2_, 0.1mM CaCl_2_). Incubation with mouse HA (1:500, Santa Cruz Biotechnology), rabbit FLAG (1:200, Cell Signaling Technology), and goat Myc (1:200, Novus Biologicals), or mouse Flag M2 (1:1000, Sigma) or mouse GFP (1:1000, Sigma) was done overnight at 4°C. Incubation with donkey anti mouse AlexaFluor647, donkey anti rabbit AlexaFluor555, and donkey anti goat AlexaFluor 488 secondary (1:500) antibodies or goat anti mouse AlexaFluor488 was done for 1hr at room temperature in the dark. Coverslips were mounted using ProLong Gold with DAPI (Invitrogen, P36931). Images were taken using an inverted Olympus IX71 microscope and processed using cellSens Dimension imaging software.

### RNA synthesis

RNA was synthesized as previously described(26). Briefly, FMR1, C9orf72, and HTT plasmid DNA was linearized with PspOMI restriction enzyme. Firefly luciferase plasmid DNA was linearized with PmeI restriction enzyme. Linearized DNAs were *in vitro* transcribed using HiScribe T7 High Yield RNA Synthesis kit (NEB) with 3’-O-Me-m7 GpppG anti-reverse cap analog (ARCA) or ApppG cap (NEB). T7 reactions were done at 37°C for 2hrs followed by DNAse treatment and polyadenylation. Synthesized RNAs were cleaned and concentrated using an RNA Clean and Concentrator-25 kit (Zymo Research) according to the manufacturer’s protocol. The quality of the RNA was determined on a denaturing RNA gel.

### *in vitro* Translation

Above described transcribed RNAs were *in vitro* translated with Flexi Rabbit Reticulocyte Lysate System (Promega) as previously described (19, 22). Briefly, translation reactions were programmed with 3nM (NL assays) or 50ng (immunoblotting) FMR1, C9orf72, and HTT mRNAs for 30min at 30°C and then incubated on ice to terminate the reaction. For NL assays, completed reactions were diluted 1:6 in Glo Lysis Buffer (Promega) followed by NanoLuc luciferase assays as described above. For immunoblotting, reactions were diluted in 6x SDS sample loading dye and boiled at 70°C for 15min.

Proteins were separated on 12% SDS-PAGE gels as described above.

### PCR for splicing products

RNA isolated from HEK293 cells transfected with different HTT NL DNA reporter plasmids was reverse transcribed using iScript cDNA synthesis kit (BioRad, 1708890) according to the manufacturer’s protocol using 1μg as input RNA. 100ng of cDNA was amplified using Taq 2x Master Mix (Fortis). Primer P1 (5’- ATTAATTGTTGCCGGGAAGCT-3’) or P3 (5’- TCCTCCGATCGTTGTCAGAA-3’) in the ampicillin resistance gene of pcDNA3.1 was paired with P2 (5’- CTTTGGATCGGAGTTACGGAC-3’) in the NL sequence. A primer was designed at the 5’ end of NL P4 (5’- TCACACTCGAAGATTTCGTTG-3’) to amplify NL alone as a control for the cDNAs. The cycling conditions were 95°C (2min) followed by 30 cycles of 95°C (30s), 50°C (30s), 72°C (2min), with a final 10min extension at 72°C. PCR reactions were run on 1.5% agarose gels.

### RNA sequencing

Total RNA was isolated from each well of a 12 well dish of HEK293 or 293T cells transfected with the different glutamine and alanine frame RAN or WT HTT reporters using the RNeasy Mini Kit from Qiagen according to the manufacturer’s protocol. Samples were DNAse treated using Qiagen’s RNase-free DNase set according to their protocol. RNA quality was assessed with the Qubit RNA IQ assay (Thermo Fisher). RNA sequencing was performed by Azenta by 2x150 bp paired end sequencing at 20- 30 million reads per sample on an Illumina sequencer.

To identify repeat ends the SATCFinder pipeline was utilized as described previously (29) and available on GITHub (https://github.com/AnkurJainLab/SATCfinder). Briefly, this module fetches reads containing a specified repeat sequence then computationally trims these reads to facilitate mapping of the non-repetitive portion of reads. This facilitates localization of “CAG ends” which are identified by the included SATCFinder ends module. This module was used with the following parameters: bbduk (BBTools version 39.10)(51) was run with the following options. Sequencing adapters and quality trimming was then performed using cutadapt (Martin et al, 2011) version 4.9 with flags “-a AGATCGGAAGAG --error-rate=0.1--times=1 --overlap=5 --minimum-length=20 --quality-cutoff=20”.

These trimmed reads were then processed using the SATCFinder python script to convert FASTQ files to a SAM file with read mates paired (33). This SAM file was then mapped to the human hg38 genome with GRCh GTF annotation file 38.109 using the STAR aligner version (39) 2.7.11b using flags “-- outSAMtype BAM SortedByCoordinate --outSAMunmapped Within –genomeFastaFiles construct.fa, readFilesType SAM PE”, with construct.fa referring to the relevant plasmid fasta file allow mapping to the transfected plasmid. The resulting BAM file was indexed using SAMtools version 1.21(39,40) and read mates separated and stored in a BAM file only containing reads with trimmed repeats. This selected BAM file was then processed with the SATCfinder ends module (33), to identify repeat ends at specific genomic coordinates. This outputs a .tsv file that allows further analysis and visualization using GraphPad Prism to generate bar plots shown in Figure 4 and Supplemental Figure 4. This .tsv file could then be used to calculate percentage of repeat ends originating at the repeat sequence as those that map within 10 base pairs of the repeat sequence divided by the total number of repeat ends. Those originating more than 10 base pairs from the repeat sequence represented reads potentially arising from splice events. Since none observed were represented by more than 100 reads, a threshold set by Anderson et al, these were not further investigated.

**Figure 2:**
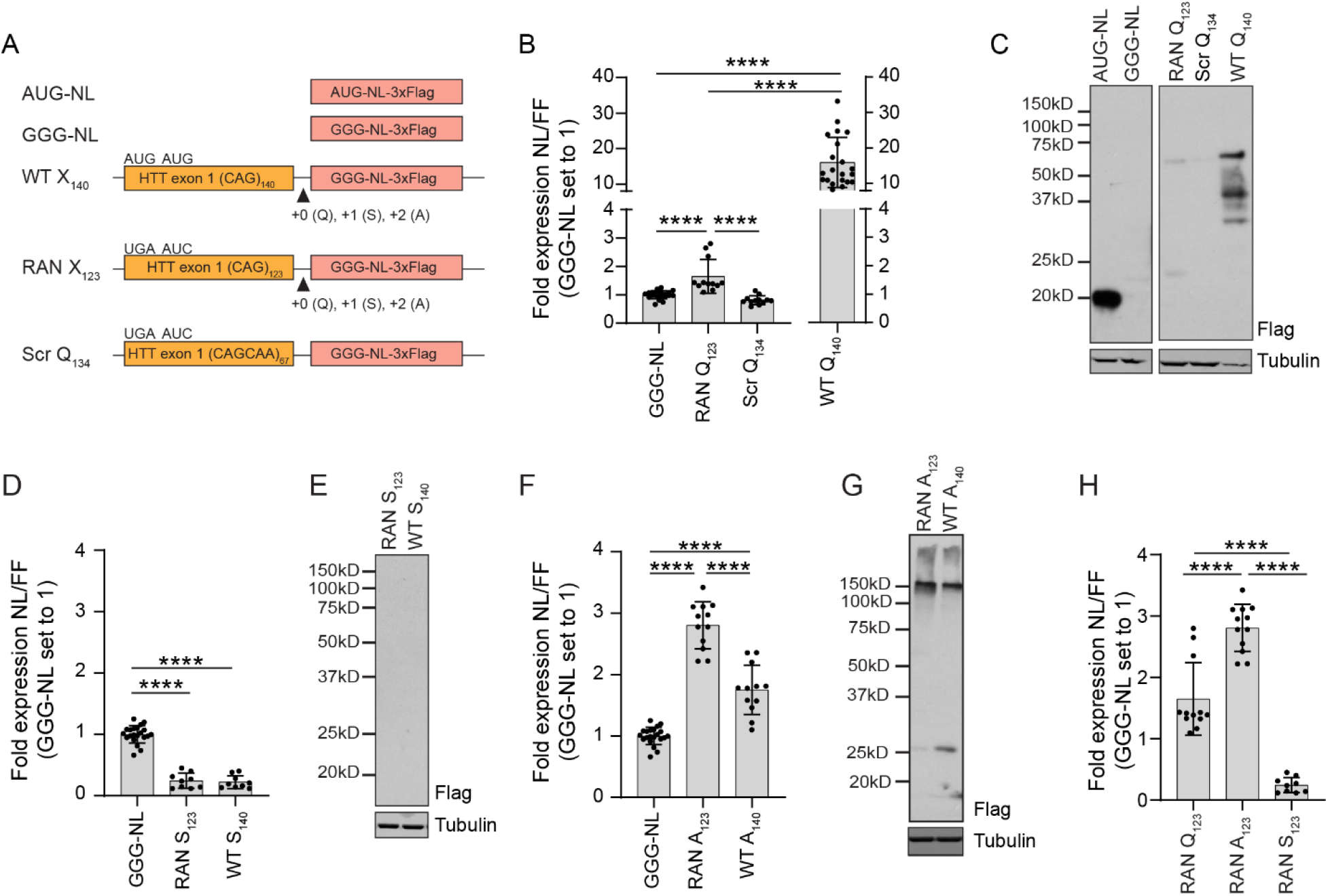
HTT RAN translation is most efficient in the alanine reading frame. A. Schematic of control and HTT reporters with NanoLuc (NL) luciferase 3xFLAG tags. B. NL assay of RAN translation in the glutamine frame from HEK293 cells transfected with glutamine DNA plasmids. Firefly (FF) luciferase was an internal normalization control. C. Representative western blot of glutamine transfected HEK293 cell lysates probed for FLAG. Tubulin was used as a loading control. D. Serine frame RAN translation efficiency from DNA transfected HEK293 cells. E. Representative western blot of HEK293 cells transfected with serine plasmids and probed with FLAG antibody. F. RAN translation of alanine frame by NL assay from transfected HEK293 cells. G. Representative western blot from HEK293 cells expressing alanine frame NL DNA plasmids. H. Comparison of RAN translation efficiencies between HTT frames. Graphs are means ± SD from triplicate samples and at least 3 different experiments (N>9). **** p<0.0001. Ordinary one-way ANOVA with Dunnett’s correction for multiple comparisons.

**Figure 3.**
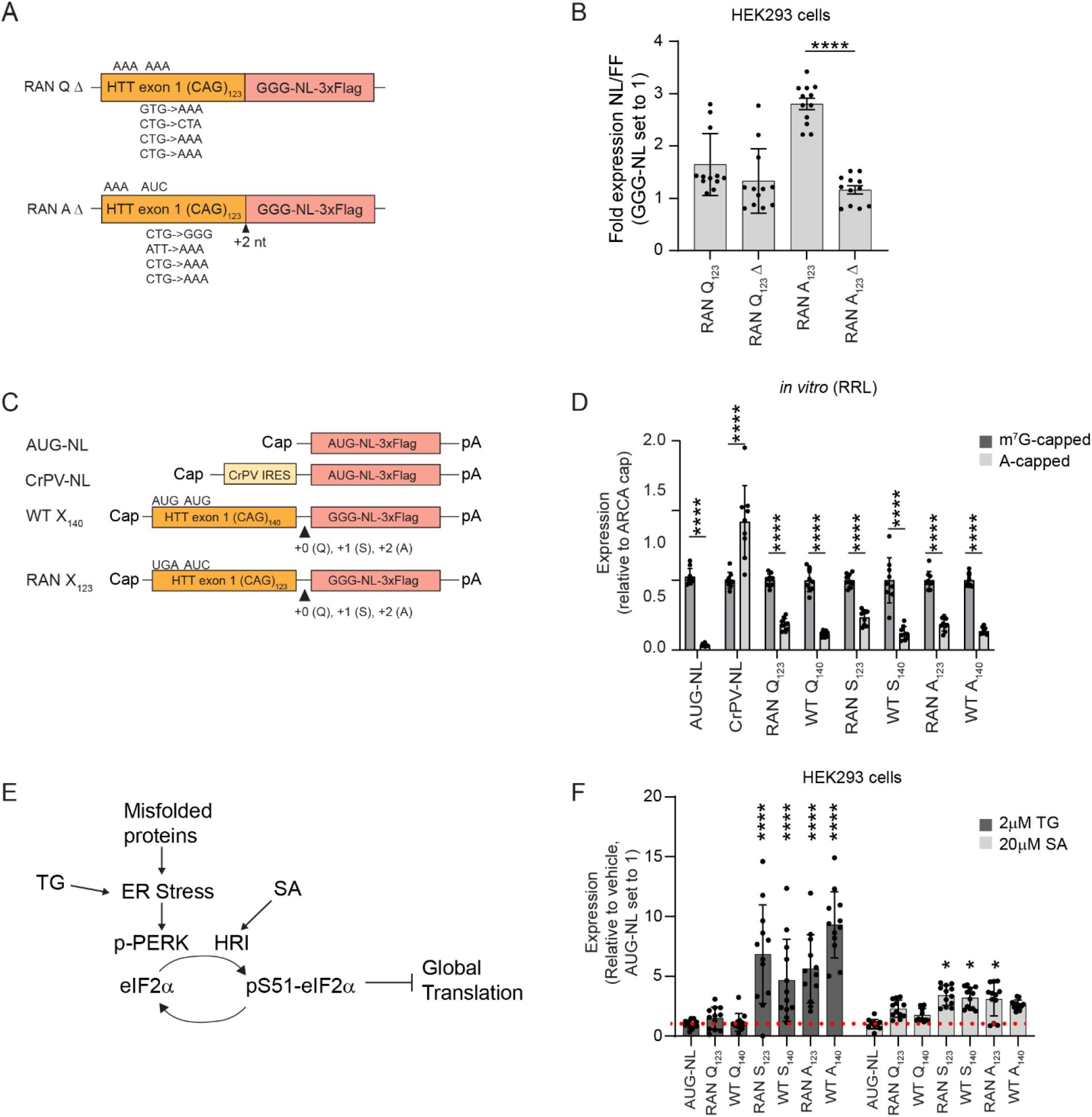
HTT RAN translation is cap-dependent and enhanced by cellular stress. A. Schematic of reporters with near-cognate codons mutated to AAA or GGG. B. NL assay from HEK293 cells transfected with near-cognate mutant plasmid DNA. FF luciferase was an internal normalization control. Comparisons are to GGG-NL control. C. Schematic of control and HTT in vitro transcribed RNAs with m^7^G (ARCA) cap or A-cap and polyA tail. D. Expression of m^7^G capped and A-capped control and HTT reporters in an *in vitro* rabbit reticulocyte (RRL) system. E. Schematic of the integrate stress response (ISR) pathway with ties of action for thapsigargin (TG) and sodium arsenite (SA). F. HEK293 cells transfected with reporter DNA and activation of the stress response using either 2μM TG or 20μM SA. Graphs are means ± SD from triplicate samples across at least 3 different experiments (N>9/sample). B: One-way ANOVA with Dunnett’s correction for multiple comparisons. D: Two-way ANOVA with Sidak’s correction for multiple comparisons. F: Welch’s student t-test. * p<0.05, ** p<0.01, *** p<0.001, **** p<0.0001.

**Figure 4.**
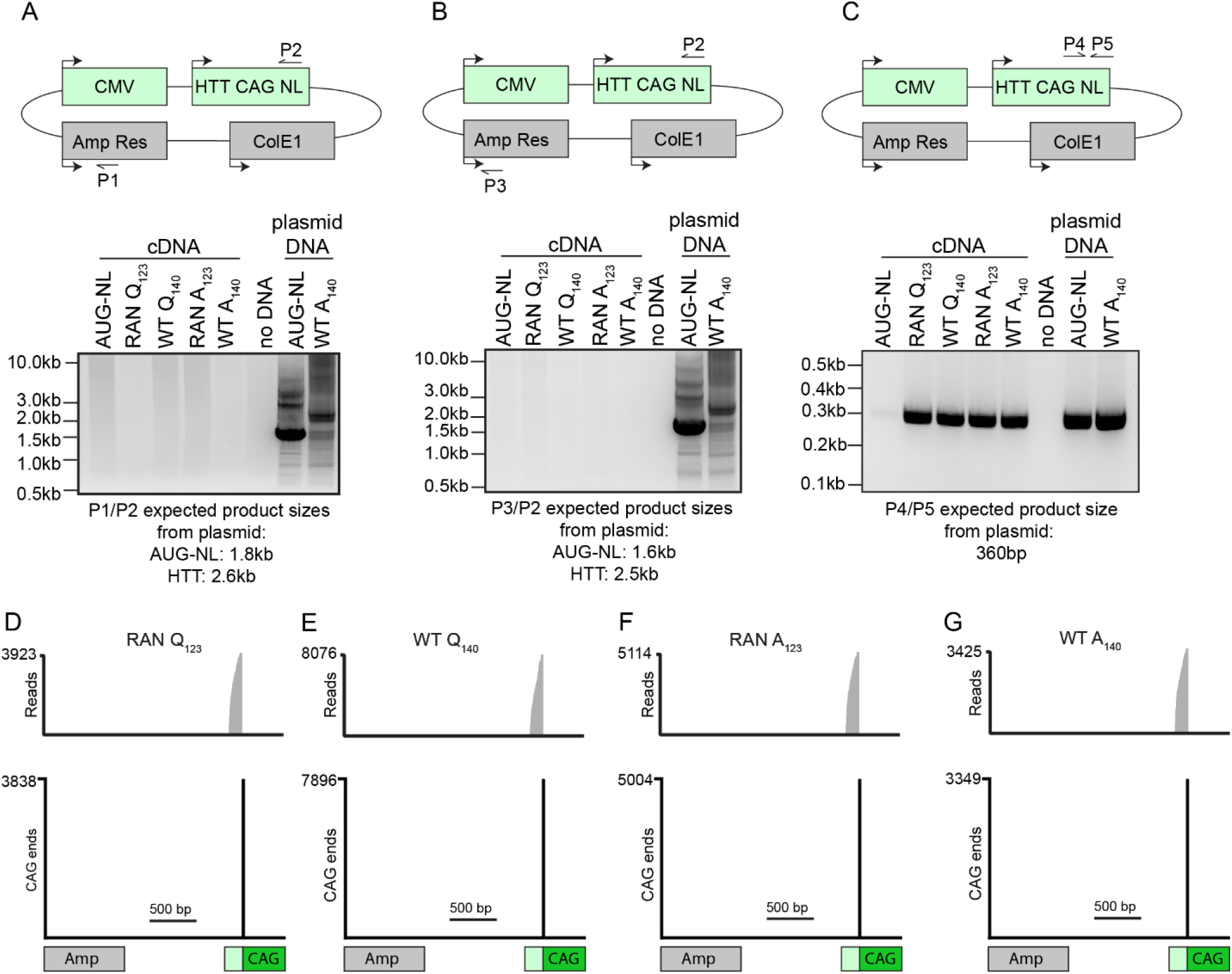
No detection of plasmid-derived mis-splicing of HTT CAG repeat-containing mRNAs. A-. C. Schematic of primer annealing sites in the HTT plasmid. PCR of cDNA reverse transcribed from RNA isolated from transfected cells does not show any amplification products. D-G. RNA-seq data looking at transcripts found in HEK293 cells transfected with HTT plasmids showed that a vast majority of the transcripts observed originated at the expected location upstream of the CAG repeat and not from a mis- splicing event.

Of note, this pipeline only analyzes reads containing the specified repeat sequence, in this case those with at least 3 consecutive CAG repeats. As the repetitive portion of are computationally trimmed before alignment, only the non-repetitive portion of reads originally containing 3 CAG repeats are successfully mapped. Two important features emerge as a result of this: (a) coverage is highest at the beginning or end of a CAG repeat tract – termed the “CAG ends” and (b) a sudden drop in coverage where the CAG repeat starts due to computational trimming of the CAG repeat prior to mapping.

### Quantitative real-time reverse transcription PCR (qRT-PCR)

RNA from transfected HEK293 cells was isolated using RNeasy Mini Kit (Qiagen). Samples were DNase treated using Qiagen’s RNase-free DNase set during the isolation process according to the protocol from Qiagen and then cleaned using Zymo Research’s RNA Clean and Concentrator-25. cDNA was generated from 1μg of cleaned RNA using the iScript cDNA Synthesis kit (BioRad). NL (custom, ID: AP7DWMP) and GAPDH (ID: Hs99999905_m1) TaqMan primer-probe pairs were used to measure cDNA abundance.

### Statistical Analyses

Statistical analyses were performed using GraphPad Prism using statistical tests denoted in figure legends. Graphs represent mean ± SD with individual data points visualized. N for each experiment is listed in the figure legends and represents biological replicates from independent experiments.

## RESULTS

### CAG repeats support RAN translation both within and outside of AUG initiated ORFs

The native HTT locus has AUG codons at positions 1 and 8 of its coding sequence, both of which are in the glutamine reading frame. Previous studies reported RAN translation from the HTT locus in all three reading frames in the absence of any upstream AUG start codon (26). To study these phenomena, we made plasmids where we placed a different tag in each reading frame (glutamine HA, serine Myc, alanine FLAG) downstream of exon 1 of HTT (Fig 1A). The WT plasmid retained both AUG codons at positions 1 and 8 of the coding sequence of HTT while in the RAN translation plasmids had these codons mutated to UGA and AUC, respectively. As expected, due to the presence of the native AUG codons, WT plasmids generated more glutamine (HA) signal compared to signal from the RAN only construct (Fig 1B). Consistent with prior work, both WT and RAN triple tagged constructs exhibited signal in the alanine (FLAG) frame (Fig 1B). Like with other polyalanine proteins, a smear was observed and the protein accumulated at the top of the gel (9, 26). Expression of the triple tag in isolation generated a signal in the Myc reading frame that matches the size of the tag itself (Fig 1B). No AUG or near-AUG cognate codon was present in the sequence 5’ to this tag to explain this product. Despite this, and consistent with some prior work using reporters for CAG RAN translation (15,37), no serine frame (Myc) product was detected - perhaps owing to its insolubility (9,30). To verify this wasn’t an antibody issue, both the HA and Myc tags were fused to the C-terminus of GFP (Sup Fig 1A) and assessed by immunocytochemistry and western blot (Sup Fig 1B-E). The HA antibody readily detected the GFP fused HA protein by western blot and ICC (Sup Fig 1B-C). While some non-specific bands were observed with the Myc antibody on western blot, it was able to detect the GFP-Myc fusion by ICC (Sup Fig 1D) and immunoblotting (Sup Fig 1E).

Similar relative expression levels of all three reading frame proteins were observed by immunocytochemistry on transfected cells. With WT triple tagged transfected cells, approximately 50% of the HA (glutamine) positive cells were also FLAG (alanine) positive while RAN triple tag cells were doubly positive about 85% of the time. In contrast, 70% of the FLAG positive cells from WT and 90% from the RAN triple tag transfected cells were HA positive (Fig 1C-E) – suggesting that alanine production is more common in cells also generating polyglutamine proteins. In contrast, no serine signal was detected by ICC using a Myc antibody in cells that stained positive for alanine or glutamine in the setting of either the WT or RAN plasmids (Fig 1C-E).

Polyglutamine repeat expansions in *HTT* trigger formation of neuronal intranuclear inclusions both in cell culture and in patient brains. However, we detected no such inclusions when we expressed the triple tag constructs in HEK293 cells. Given past studies demonstrating such inclusions often included a GFP fusion to exon 1 of *HTT* (41–44), we swapped the triple tag for a version of GFP without an AUG start codon in each potential reading frame (Sup Fig 1F) and transfected HEK293 cells to assess protein localization and aggregate formation. Consistent with prior work, WT polyglutamine fused to GFP formed intranuclear inclusions in HEK293 cells (Sup Fig 1G) after 48hrs. In contrast, RAN generated polyglutamine was diffuse throughout the cell. Polyalanine GFP signal was also diffuse throughout HEK293 cells (Sup Fig 1G) and comparable between WT and RAN constructs. In contrast, RAN translated polyserine fused to GFP was detectable in the nucleus, which was different than what we observed using the Myc tag but was consistent with prior work (30). These data suggest that polyserine products are generated by RAN translation at low levels.

### HTT RAN translation differs between reading frames

To create a more quantitative assay for measuring RAN translation efficiency from HTT exon 1, we generated a series of HTT RAN translation specific NanoLuciferase (NL) luciferase reporters by replacing the triple tag with a mutated version of NL where the AUG start codon of the NL protein was mutated to GGG (Fig 2A) (22,26). In addition, a 3x FLAG tag was included at the end of NL to aid detection by immunoblot. Consistent with prior work,(22,26) the mutant form of NL absent any upstream repeat sequence generates little signal (∼200-fold difference) compared to its AUG-NL counterpart (Sup Fig 2A). To make sure any translation we see is not due to differences in transcription of the plasmids, we did qRT-PCR to assess the RNA levels of each plasmid transfected into HEK293 cells. Transcription of the FMRpolyG reporter was elevated compared to AUG-NL, as expected(45,46), but the levels of transcription of HTT plasmids were not significantly different than AUG-NL (Sup Fig 2B). Immunostaining of NL-tagged reporters was similar to staining observed with triple tag and GFP reporters: glutamine and alanine frame reporters showed diffuse staining (Sup Fig 2C). Serine frame reporters were still lowly expressed but nuclear puncta were observed like what was seen with the GFP tag. Driving expression by inserting an upstream AUG codon in the serine frame increased the number and size of these puncta (Sup Fig 2C).

RAN translation was relatively inefficient in the glutamine frame relative to AUG-initiated translation (7%) but was still significantly greater than a negative control by NL assay (Fig 2B). Consistent with prior studies, replacement of an expanded pure CAG repeat with a CAGCAA repeat (Scr Q) that still encodes glutamine but doesn’t readily form a hairpin structure decreased translation when compared to a pure CAG repeat (Fig 2B)(9,31). A similar pattern was observed when these proteins were visualized by western blot (Fig 2C).

To quantify RAN translation in the serine and alanine reading frames, we inserted one or two nucleotides between the repeat and GGG-NL to shift the HTT reading frame to the serine or alanine reading frames, respectively. When RAN translation of the serine reading frame was assessed by NL assay, no signal was detected above the level of our GGG-NL negative control (Fig 2D) and not observed on a western blot (Fig 2E). Because the serine product may be insoluble, we collected cytoplasmic, nuclear, and insoluble fractions from transfected cells. Serine reading frame RAN translation from either 20 or 123 repeats was not observed in any fraction (Sup Fig 2D). In contrast, driving expression through the repeat with an AUG codon in the serine reading frame allowed for visualization of constructs with 20 CAG repeats in all fractions while the AUG initiated 123 CAG repeat polyserine protein was only observed in the insoluble fraction.

In contrast, RAN translation of expanded alanine repeats was significantly higher than our control construct (Fig 2F) and higher than RAN translation seen in the glutamine reading frame (Fig 2H). On western blot, expanded repeat expressing proteins accumulated at the top of the blot (Fig 2G), as was previously observed with CGG and CCG alanine repeat proteins fused to NL or GFP(13).

### Stress enhances HTT RAN translation – but not AUG initiated HTT translation

Hallmarks of RAN translation at CGG repeats in *FMR1* and GGGGCC repeats in *C9orf72* associated FTD and ALS are the use of near-cognate codons for initiation in at least some reading frames, 5’ m^7^G cap dependence in cell lines and in vitro translation assays and altered expression in response to activation of cellular stress pathways (22,24,26,29,47,48). To assess the use of near-cognate codons in the context of CAG repeats in HTT exon 1, we generated different mutants in the glutamine and alanine frames (Fig 3A & Sup Fig 3A). The HTT gene has two AUG codons at the beginning of the coding sequence. When the first AUG was mutated to a UAA stop codon and the second AUG remained a start codon, translation in the glutamine frame was intermediate to levels of WT translation and RAN translation (Sup Fig 3B). In the alanine frame, the presence of either AUG codon decreased NL compared to the RAN alanine reporter (Sup Fig 3C).

In our RAN reporter constructs, the second AUG was initially mutated to AUC, a near-cognate codon. To determine if this near cognate codon contributed to RAN translation of either glutamine or alanine, this codon was mutated to an ACC. This mutation did not change the levels of RAN translation observed in either the glutamine or alanine frames (Sup Fig 3D-E). Removing all the near-cognate codons did not change glutamine RAN translation via NL assay (Fig 3B). In contrast, mutation of near-cognate codons in the alanine frame did significantly reduce the amount of translation observed by NL assay (Fig 3B). In the alanine frame, there is one near-cognate CUG codon in the HTT 5’UTR in a reasonable Kozak context. When this site was mutated in isolation, we observed a similar decrease in NL signal as when all sites were mutated (Sup Fig 3F) – suggesting a functional role of this CUG in HTT RAN initiation.

We *in vitro* transcribed RNA with a functional m^7^G (ARCA) cap or a non-functional A-cap to determine if RAN HTT translation depends on 5’ m^7^G cap binding (Fig 3C). In an *in vitro* translation rabbit reticulocyte lysate system, A-capped HTT mRNAs in all 3 reading frames exhibited decreased expression compared to functional ARCA capped mRNAs (Fig 3D). However, the degree of cap dependence for these RNAs was intermediate between that of a cap-dependent AUG-NL mRNA and that of a CrPV-NL positive control for cap-independent translation. As expected, WT glutamine was more cap dependent (Fig 3D) than the RAN glutamine mRNA. Raw NL values are shown in Sup Fig 3G.

Another aspect of RAN translation is the increased translation in response to activation of the integrated stress response (ISR)(26). We treated HEK293 cells with either thapsigargin (TG, ER stress) or sodium arsenite (SA, mitochondrial dysfunction stress) to induce the integrated stress response (Fig 3E). RAN alanine was increased when the integrated stress response was activated by both stressors (Fig 3F). In contrast, neither RAN nor WT glutamine translation increased when the ISR pathway was activated by either stressor (Fig 3F). Interestingly, RAN and WT serine translation – despite having an extremely low signal at baseline - was significantly increased when the ISR was activated, suggesting that generation of the serine product *in vivo* may result from a cellular response to stress (17,31).

### RAN translation from plasmid HTT does not result solely from aberrant splicing

A recent report suggests that RAN translation from CAG repeat containing reporter constructs may be an artifact brought on by cryptic transcription initiation within the plasmid and splicing into the repeat with the repeating “AG” serving as a splice acceptor(33). To assess whether aberrant splicing from our plasmids might occur, we first designed primers to the predicted previously described mis- splicing events from the ampicillin resistance gene of pcDNA3.1 into the CAG repeat. Two different primers binding to different portions of the ampicillin gene and a single reverse primer in the NL gene were used (Fig 4A-B). A primer set was designed inside the NL sequence to act as a control for the generation of the cDNA (Fig 4C). RNA isolated from transfected cells was reverse transcribed into cDNA and the cDNA was used as the PCR template. No aberrant PCR products were observed using these primer sets (Fig 4A-B), but products were readily detected when using primers in the NL sequence (Fig 4C). Next, we performed RNA-seq on samples after transfection with plasmids expressing different RAN translation reporters. No mis-splicing events were identified (Fig 4D-G, Sup Fig 4). In contrast, we did observe previously described splicing into native CAG repeat sequences (Sup Fig 4A- B)(33). Because initial reports of mis-splicing was assessed in HEK293T cells, we repeated the RNA-seq analysis on samples from transfected HEK293T cells. No differences were observed in these cells (Sup Fig 4D-G). We conclude that in the context of our pcDNA3.1 plasmid and the HTT surround sequence in HEK293 cells, such mis-splicing events are either not present or exceedingly rare.

If this aberrant splicing event were the sole contributor to RAN translation, then *in vitro* transcribed RNAs would not be expected to support RAN translation, as they would not be subject to splicing. To test this, we *in vitro* transcribed RAN and WT CAG reporter RNAs with a functional m^7^G cap (Fig 5A) and repeated our earlier assays. Translation from IVT RNAs was generally higher than that seen with plasmids (Fig 5B). If mis-splicing were to occur, it was thought to enhance translation of the alanine frame. Translation of RAN alanine from DNA was similar to RAN translation observed from IVT RNA. Induction of the integrated stress response by sodium arsenite had similar effects on CAG RAN translation between RNA and DNA transfections (Fig 5C).

**Figure 5.**
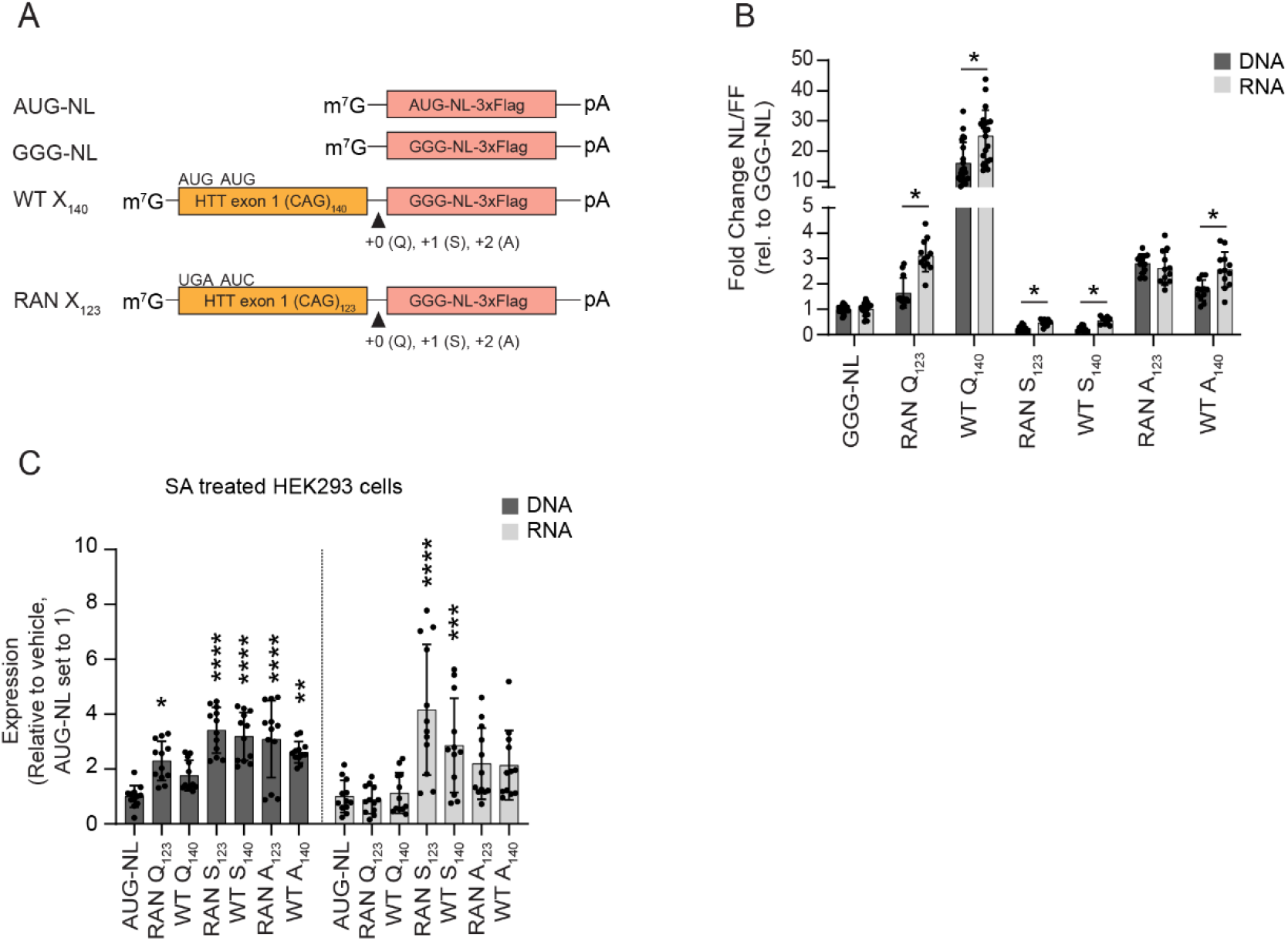
Comparing the efficiency of HTT RAN translation from DNA and RNA reporters. A. Schematic of control and HTT *in vitro* transcribed RNAs with m^7^G cap and polyA tail. B. NL assay of HEK293 cells transfected with plasmid DNA or IVT RNAs. FF luciferase was an internal normalization control. C. Comparison of 20μM sodium arsenite (SA) impact on HTT CAG RAN translation from DNA or RNA based reporters. Graphs are means ± SD from triplicate samples and at least 3 different experiments. B: Welch’s multiple t-tests. C: Two-way ANOVA with Sidak’s correction for multiple comparisons. * p<0.05, ** p<0.01, *** p<0.001, **** p<0.0001.

### RAN translation efficiency differs by repeat, reading frame, and cellular context

To assess the comparative efficiency of RAN translation from CAG repeats versus other repeats, we performed parallel assays comparing CAG repeats to existing *in vitro* transcribed RAN translation RNAs for CGG and GGGGCC repeats in an *in vitro* rabbit reticulocyte lysate translation system, transfected HEK293 cells (22,26), and rat primary hippocampal cultures (Fig 6A). We used in vitro transcribed RNA rather than plasmid transfection to avoid any concerns related to potential mis-splicing events associated with plasmids and to better control for positive and negative effects of each repeat on RNA transcription rates (Sup Fig 2B)(45,46,49). Previously described RAN translation reporters from *FMR1* and *C9orf72* in 2 reading frames were used to generate comparisons of RAN translation efficiency (Fig 6B)(26,50).

**Figure 6.**
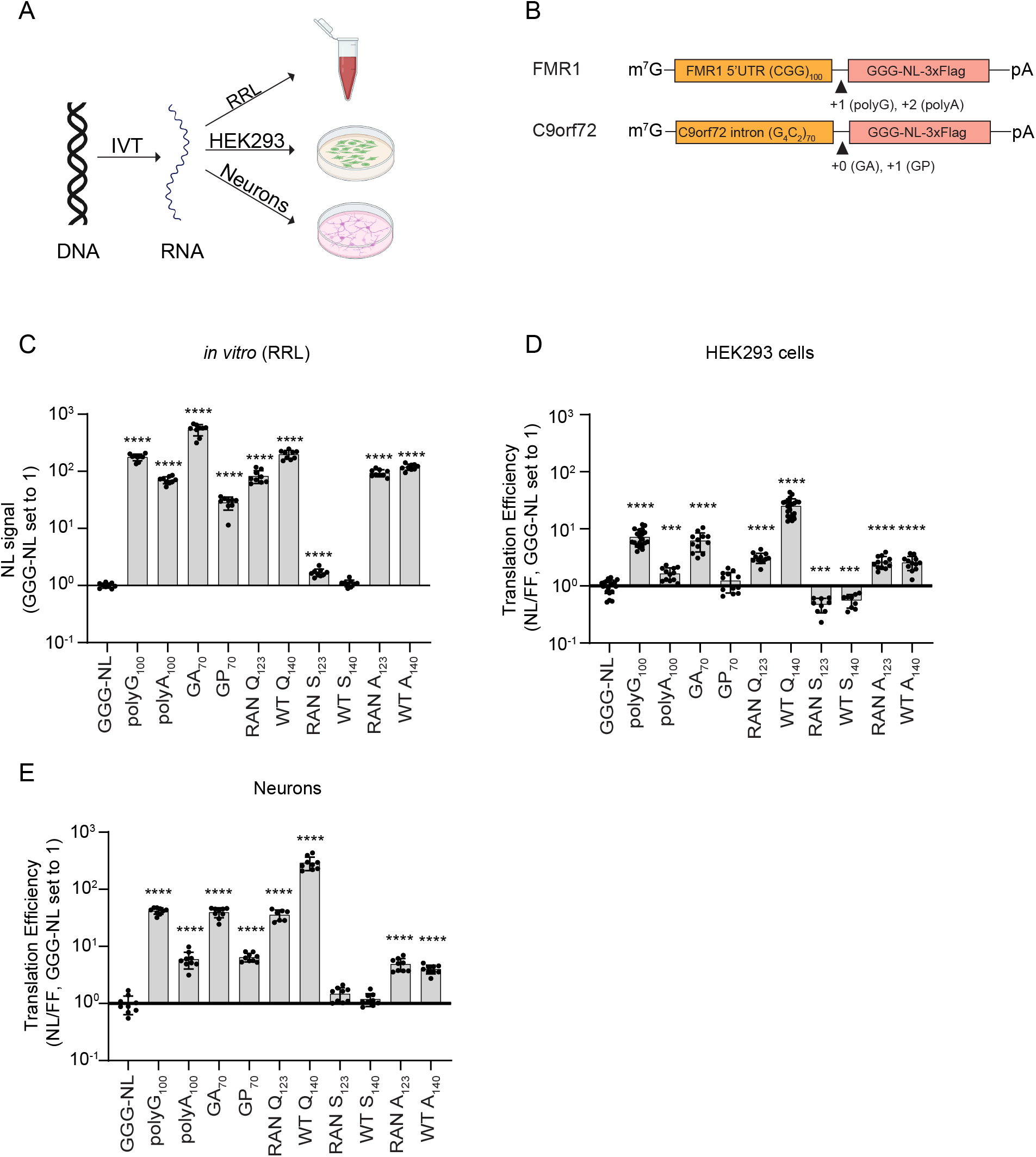
RAN translation efficiency across different repeats, reading frames, and cell types. A. Schematic of *in vitro* transcribed RNA from DNA and transfected into 3 different model systems. B. Schematic of *FMR1* and *C9or72 in vitro* transcribed RNAs with m^7^G cap and polyA tail. Comparison of RAN translation efficiencies in a rabbit reticulocyte lysate system (C), HEK293 cells (D), and rodent primary neurons (E). Graphs are means ± SD from triplicate samples and at least 3 different experiments. C-E: Welch one-way ANOVA with post-hoc Dunnet’s T3 multiple comparisons test vs GGG-NL. * p<0.05, ** p<0.01, *** p<0.001, **** p<0.0001.

In rabbit reticulocyte lysates, translation from the *C9orf72* polyGA was the highest (500x higher than GGG-NL), followed by FMRpolyG (170x). RAN glutamine (80x) and both WT and RAN alanine (100x and 120x, respectively) were similar to FMRpolyA (70x) and GP (33x) translation. Translation in the HTT serine reading frame was not higher than the negative control (Fig 6C). In HEK293 cells, RAN translation from *in vitro* transcribed RNAs was an inefficient process for all transcripts. FMRpolyG was the most efficiently RAN translated transcript (7x higher than GGG-NL), followed by GA (4x). RAN glutamine (3.2x) and RAN/WT alanine (both 2.7x) were comparable to FMRpolyA (1.6x). (Fig 6D). RAN translation signal in the serine reading frame was less than the negative control.

In rodent neurons, CAG RAN translation from IVT RNAs was significantly higher than HEK293 cells. RAN glutamine translation was similar to FMRpolyG and GA translation (all approximately 38x versus 41x and 45x higher than GGG-NL, respectively). RAN translation in the alanine reading frame was 4-5x the negative control for the assay (Fig 6E). Translation of serine was not above the background level of GGG-NL. Immunoblotting showed comparable abundance to that observed with NL reporters (Sup Fig 5). By immunoblot, IVT RAN alanine reporter RNA generated detectable products in RRL and HEK293 assay systems and inconsistently detectable products in neurons (Sup Fig 5G-I), perhaps owing to neuron’s overall lower translational efficiency.

## DISCUSSION

Expansion of short tandem repeats are associated with over 60 age-related neurodegenerative disorders. Until recently, how repeats elicit disease was largely inferred from their surrounding sequence context, with an assumption that trinucleotide repeat expansions within open reading frames would elicit toxicity predominantly through AUG initiated translation of the repeats into toxic (typically polyglutamine) products that accumulate within intraneuronal inclusions(51,52). The surprising finding that repeats sometimes support non-AUG initiated translation of proteins both outside of annotated open reading frames (as occurs in FXTAS and *C9orf72* associated FTD/ALS) as well as in different reading frames of repeats within annotated open reading frames (as occurs at CAG repeats in the *HTT* locus) creates a complicated scenario whereby multiple potential toxic protein species may synergistically contribute to toxicity (53). Here we studied how “in-ORF” RAN translation events at CAG repeats in the *HTT* sequence context occur and compared it to RAN translation from CGG and GGGGCC repeats embedded within a 5’ leader sequence outside of an ORF and assessed their different properties and requirements. RAN translation from CAG repeats in the glutamine and alanine reading frames are detectable from both plasmid based reporters and from *in vitro* transcribed RNA reporters in both cell lysate based systems and transfected neurons with only modest impacts regarding the presence of upstream out-of-frame AUG codons (as happens natively in exon1 of HTT) - largely consistent with prior studies of CAG repeats in SCA8 and previous mechanistic studies of this repeat within the HTT, Ataxin-2 and Ataxin-3 sequence contexts (9,15,16,30). RAN initiation and production from CAG repeats behaved similarly to that previously reported for CGG and GGGGCC repeats, exhibiting cap-dependence in a reticulocyte lysate and enhancement by activation of the integrated stress response in human cell lines (24,26,47,54).

These data also largely support the original finding that CAG repeats trigger translational initiation absent an AUG start codon(9) and that products from this initiation cannot be fully explained by a plasmid-based technical artifact or alternative splicing event (33,55).

We observe different efficiencies for RAN translation from different repeats across different sequence contexts (as assessed in reporter assays). Prior to this study, data generated by our group and others suggested that RAN translational initiation is significantly more efficient from near-cognate initiation codons just outside of the repeats (as occurs at an ACG codon in the +1 FMRpolyG reading frame of *FMR1* CGG repeats or a CUG codon in the GA reading frame of GGGGCC *C9orf72* repeats) than it is from non-AUG cognate initiation events within the repeat itself (as is thought to occur in the +2 Alanine reading frame of CGG repeats in *FMR1* and the GP reading frame of GGGGCC *C9orf72* repeats) (22,24,26,29). However, our findings suggest that RAN translation from the glutamine reading frame of CAG reporter constructs occurs quite efficiently in neurons – on a level comparable to GA DPR production (Fig 6E) in *C9orf72* – despite the absence of any near-AUG cognate codon available for use in initiation. In contrast, at least some of the translation observed in the alanine reading frame does use a near-cognate CUG codon located in the 5’UTR of HTT. Some of these observed effects may be explained by differences in the rates of elongation through different repeat sequences post-initiation, as G-rich sequences and arginine containing open reading frames are more likely to generate ribosomal stalls (56,57). Alternatively, CAG repeats may create a sequence which is uniquely capable of engaging ribosomes and supporting initiation. What seems less likely to provide an explanation here is ribosomal frameshifts from one reading frame into another as others have proposed (56–63), because mutation of AUG codons in the polyglutamine reading frame had no impact on HTT RAN translation in the polyalanine reading frame from in vitro transcribed RNA (Fig 5B).

An alternative explanation for how polyalanine products might be generated was provided by work from Anderson et al, who observed the generation of cryptic RNAs from pcDNA3.1 plasmids that used the CAG repeat as a splice acceptor(33). Their work suggested that much of the signal observed in the polyalanine reading frame of the CAG SCA8 RAN reporters used in the original manuscript describing RAN translation emerged from this abnormal splice event bringing an AUG codon in-frame with GCA repeats to make this product. Our own data provides mixed support for this hypothesis. IVT generated CAG repeat RNAs still supported RAN translation in an *in vitro* RRL lysate, in HEK293 cells, and in transfected rodent neurons - albeit with lower efficiency in some reading frames and in some cellular contexts (Fig 6). We did not observe the mis-spliced RNA products that were previously reported from CAG SCA8 RAN reporters in our transfection studies by either directed RT-PCR or by RNA-seq in either HEK293 or HEK293T cells, despite observing thousands of plasmid derived repeat-containing reads in our samples (Figure 4F-G and Sup Fig 4F-G). As such, our data suggest that aberrantly spliced templates are unlikely to explain the totality of our findings or those of others in the field.

Using a variety of reporter systems, we were unable to reliably detect polyserine product signal from the CAG repeat in HTT exon 1 at levels above the background level of detection in our assay, which is consistent with some (15,16) but not all (9,17,30) prior studies. This may reflect the insolubility of polyserine(30). Consistent with this idea, we did observe polyserine signal by ICC when the HTT exon 1 was fused to GFP (Sup Fig 1G). However, we did not observe it within the insoluble fraction by immunoblot when CAG repeats were fused to NL (Sup Fig 2C). While we believe that technical reasons explain this discrepancy, RAN translated proteins were consistently generated less efficiently in this reading frame across systems in our hands. Despite this, as with RAN translation of CGG and GGGGCC repeats, RAN translation of both polyalanine and polyserine products from the CAG repeat in HTT exon 1 were enhanced by activation of the integrated stress response (Fig 3F, Sup Fig 5C). However, neither AUG initiated nor RAN initiated translation of polyglutamine proteins exhibited this same response. This finding is particularly intriguing based on evidence that both polyserine and polyalanine products accumulate at later stages of disease in HTT BAC transgenic mice with pure CAG repeats and in CAG repeat expansion disorder patients, but not in mice with interrupted CAGCAA repeats (31). Accumulation of these products may reflect a late event in polyglutamine-induced neurodegeneration that triggers activation of the ISR and favors production of these protein products (17).

This manuscript has a number of limitations. First, it relies on reporter constructs which are not expressed from the native human sequence locus with relatively limited CAG repeat sizes. While these tools allow for greater fidelity in measurement and biochemical assessments, they may not fully recapitulate the behavior of all of the endogenous transcripts derived from this disease associated loci. In particular, recent studies suggesting massive somatic instability in disease tissues raise the possibility that large expansions may exhibit different patterns of behavior in terms of aberrant splicing, translational initiation, and product generation. Second, while we do not observe any aberrant RNA products generated from our plasmids, it is possible that deeper sequencing might reveal rare events consistent with those observed previously. Even rare mis-splicing events could generate efficient templates for translation. Thus, while our in vitro transcribed RNA and plasmid-based reporters exhibit largely similar findings, a contribution of such aberrant products to DNA based reporter assays cannot be ruled out.

Lastly, it remains unclear what contribution CAG derived RAN translated proteins might have on pathogenesis in Huntington Disease. Future studies will be needed to delineate whether their production is a relevant therapeutic target.

In summary, we observe that CAG repeats in the context of exon 1 of *HTT* can support RAN translation and that this translation shares many key features with RAN translation at CGG and GGGGCC repeats. Moreover, our data suggests that RAN translation from CAG repeats is unlikely to result solely from a plasmid splicing artifact. As we define the mechanisms of RAN translation at different repeats, we will hopefully reveal unique and shared targetable pathways that have the potential to mitigate neurodegeneration in Huntington disease as well as the many other currently untreatable nucleotide repeat expansion disorders.

## ACKNOWLEDGEMENTS

The authors would like to thank Dr. William Yang for sharing HTT plasmids. We also thank former and current Todd lab members for discussions and technical advice.

## AUTHOR CONTRIBUTIONS

AK and PKT conceived the project. AK, RMA, and CJM performed experiments. RMA analyzed the RNA-seq data set. AK wrote the initial draft of the manuscript with input from RMA, CJM, and PKT. All authors reviewed and edited the manuscript.

## DATA AVAILABILITY

RNASeq data are available through GEO (accession number 25440016). All other study data are available in the article and/or the supplementary information.

---

**Supplemental Figure 1.**
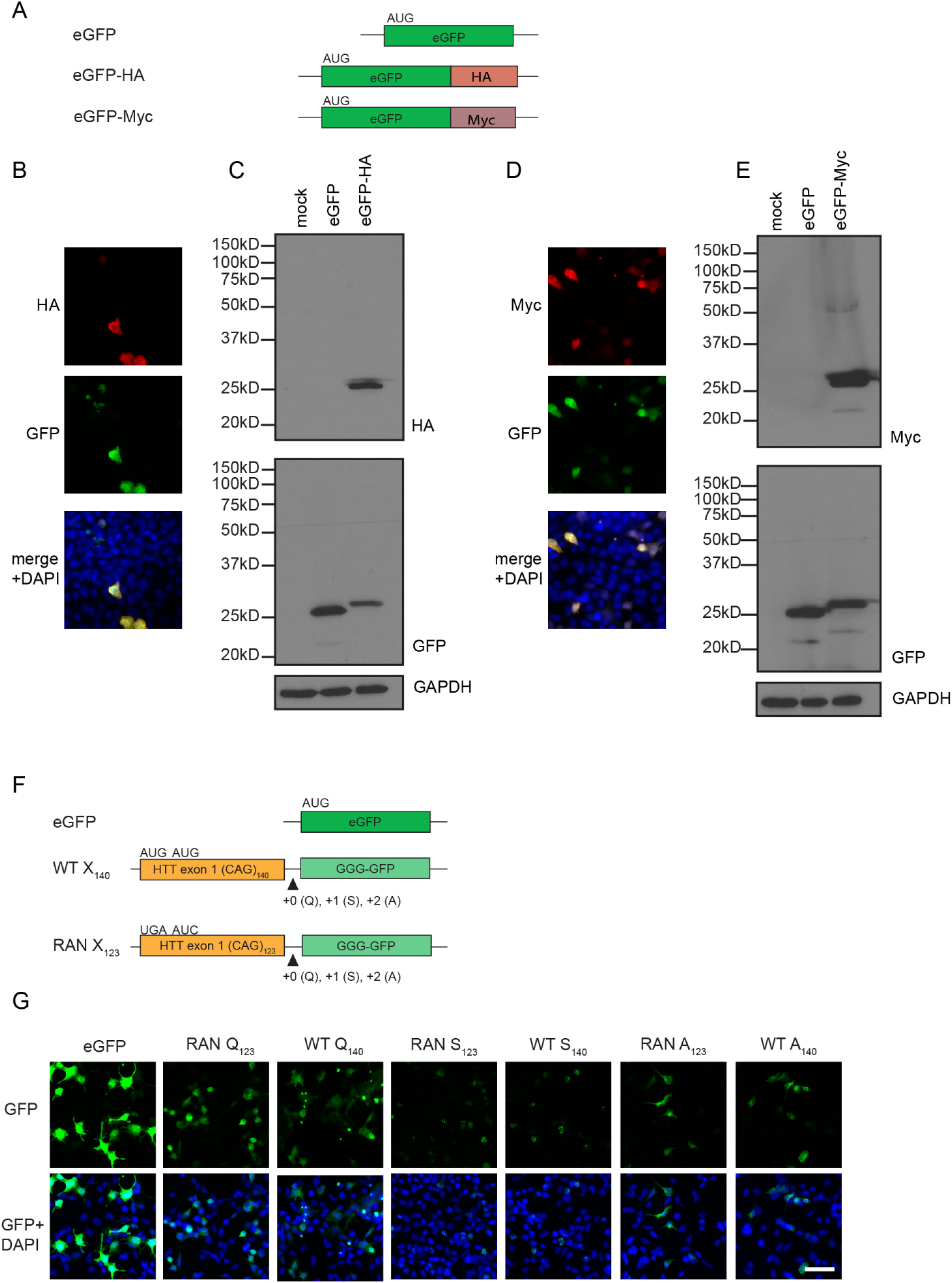
Localization of RAN translation products from HTT reporters. A. Schematic of the eGFP control plasmid and eGFP plasmids with either a C-terminal HA or Myc tag. Representative images and western blots of HA (B-C) and Myc (D-E) in HEK293 cells transfected with eGFP with a C-terminal HA or C-terminal Myc tag. GAPDH was used as a loading control. F. Schematics of HTT plasmids with a C-terminal eGFP without an AUG start codon. G. Representative images of HEK293 cells 48hrs after transfection with the indicated GFP (G). DAPI was a counterstain for nuclei. Scale bar is 50μm.

**Supplemental Figure 2.**
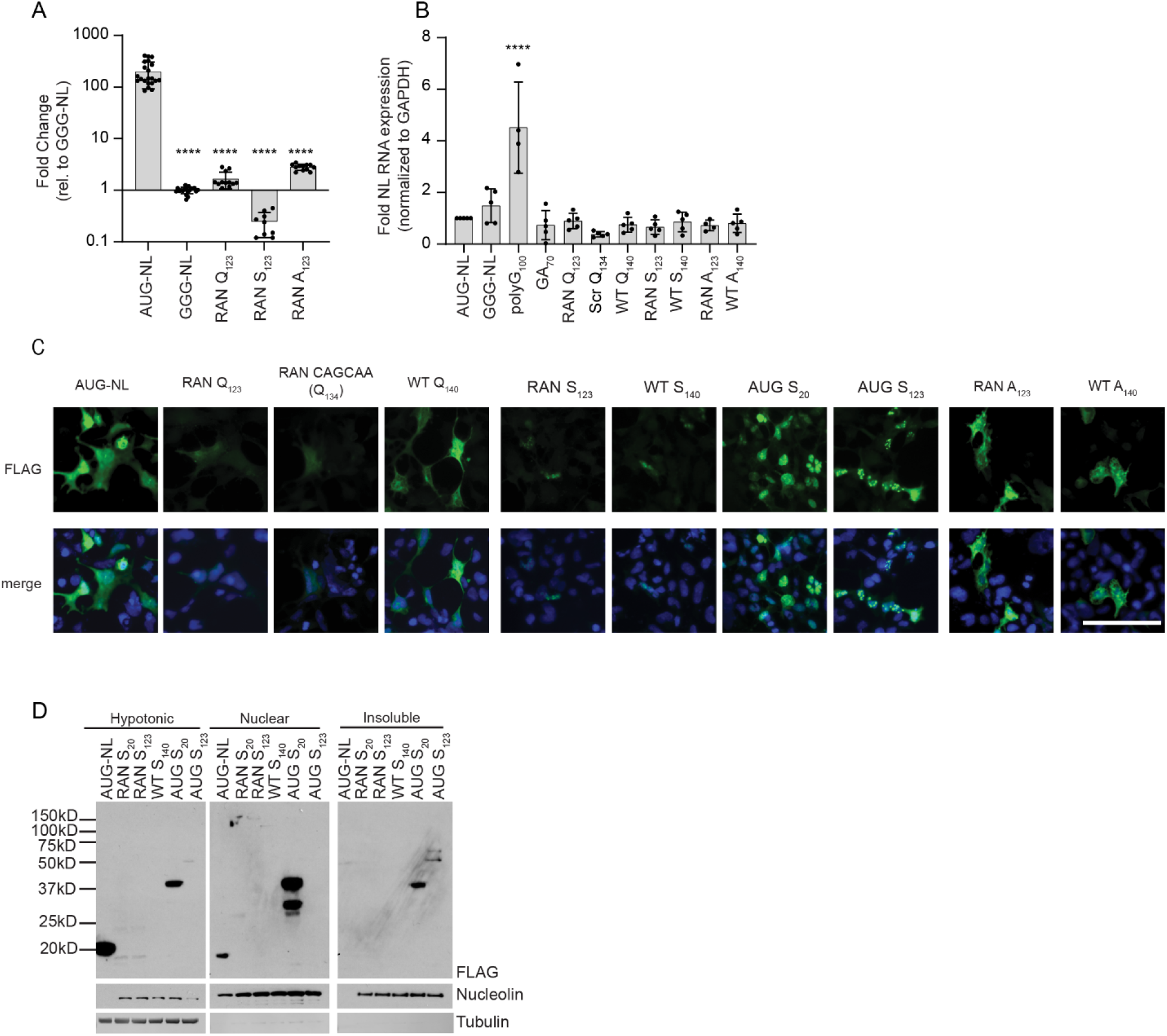
RNA and protein expression from different RAN translation reporters. A. NL assay comparing luminescence of HTT RAN expressing plasmids to AUG- or GGG-NL. B. Relative mRNA expression for identified constructs in transfected HEK293 cells. GAPDH was used as a normalization control. C. Immunocytochemistry of NL reporters. D. Representative western blot of polyserine products in different cellular fractions of transfected HEK293 cells. Nucleolin was used as a nuclear marker. Tubulin was used as a cytoplasmic/hypotonic marker. Graphs are means ± SD from triplicate samples and at least 3 different experiments. A-B was an ordinary one-way ANOVA with Dunnett’s correction for multiple comparisons. **** p<0.0001. Scale bar in C is 50μm.

**Supplemental Figure 3.**
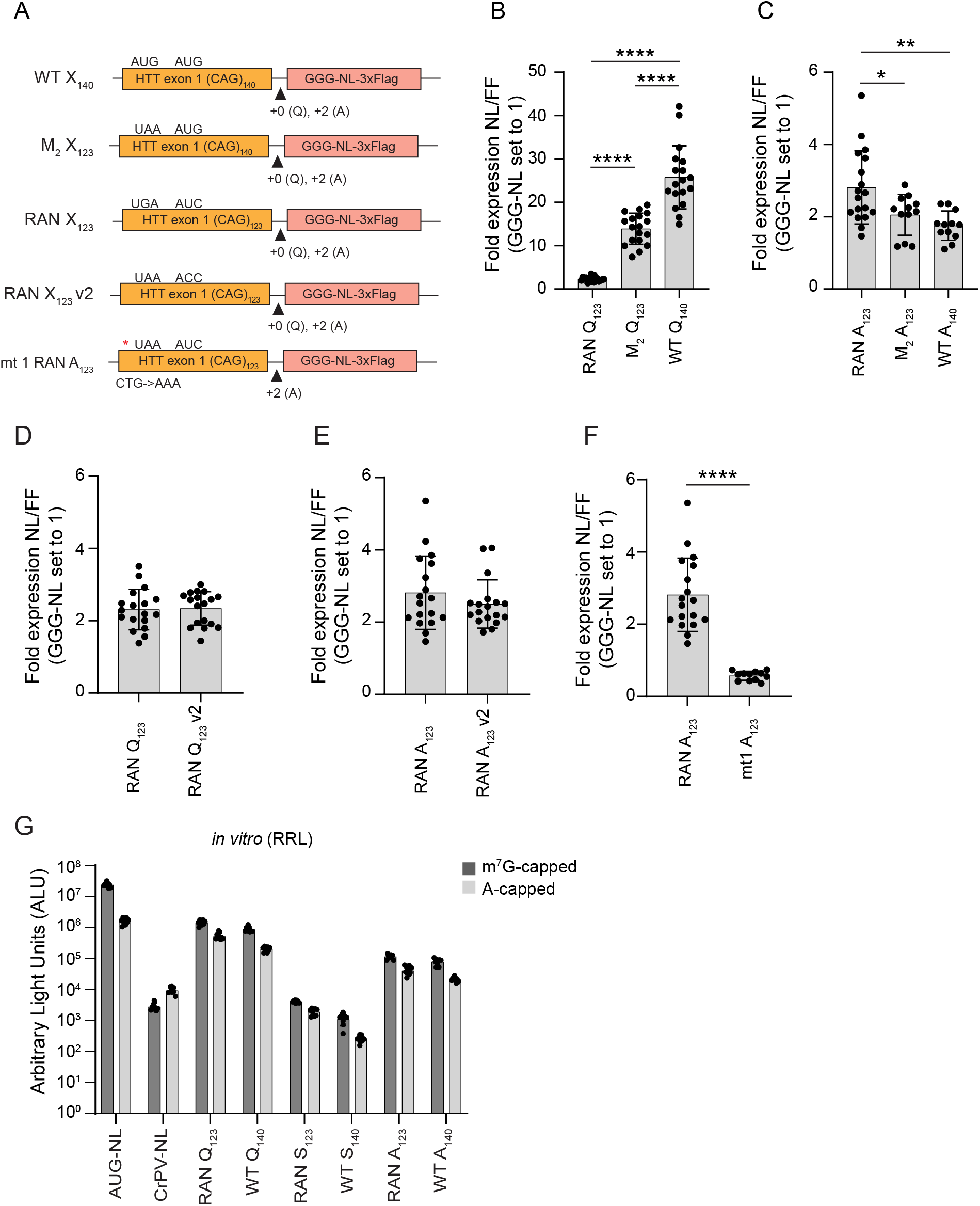
Individual mutation analysis with RAN translation HTT reporter plasmids. A. Schematic of the different codon mutations made to either glutamine or alanine frame HTT plasmids. * represents the location of the mutated CUG codon. B-C. NL assay comparing the effect of the second M codon on glutamine translation or alanine translation, respectively. D-E. NL assay comparing the effect of mutating the second M to a non-near-cognate codon on the translation of the glutamine or alanine frame, respectively. F. NL assay of the effect of the near-cognate CUG codon on alanine RAN translation. G. Raw NL values of m^7^G-capped and A-capped mRNAs in the RRL system. Graphs are means ± SD from triplicate samples and at least 3 different experiments. B-C was an ordinary one-way ANOVA with Tukey’s correction for multiple comparisons. D-F was Welch’s t-test. * p<0.05, ** p<0.01, *** p<0.0001.

**Supplemental Figure 4:**
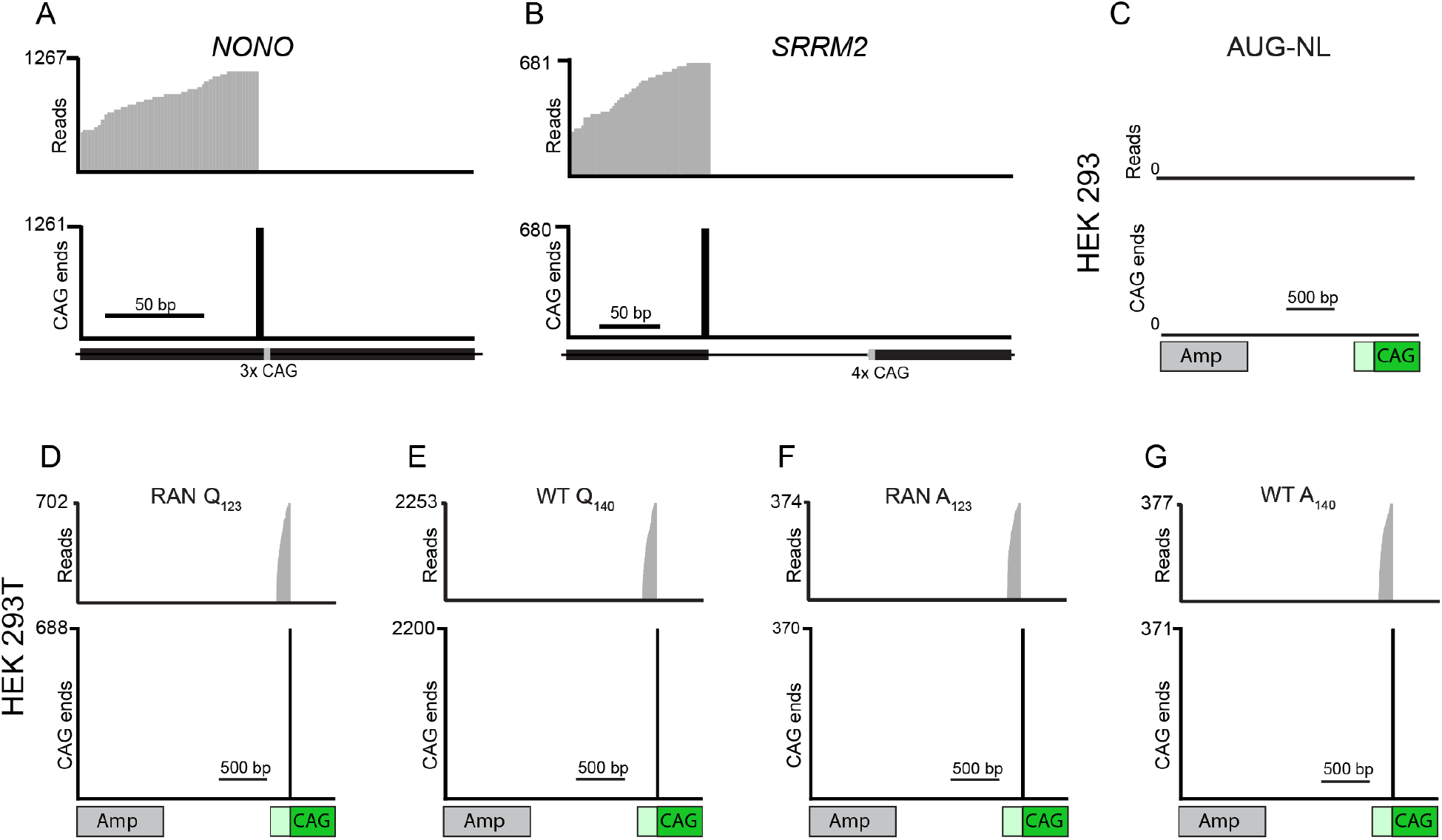
Assessment for 5’ ends of CAG repeat containing transcripts derived from genomic genes and from additional plasmids in HEK293 and HEK293T cells. CAG ends determined using the SATCFinder pipeline for the genes *NONO* (A) and *SRRM2* (B) for HEK293 cells transfected with the AUG-NL plasmid. CAG ends in the transfected plasmid in HEK293 cells for the AUG-NL (C). CAG ends for the transfected plasmid in HEK293T cells for the RAN Q_123_ (D), WT Q_140_ (E), RAN A_123_ (F), and WT A_140_ (G) plasmids.

**Supplemental Figure 5:**
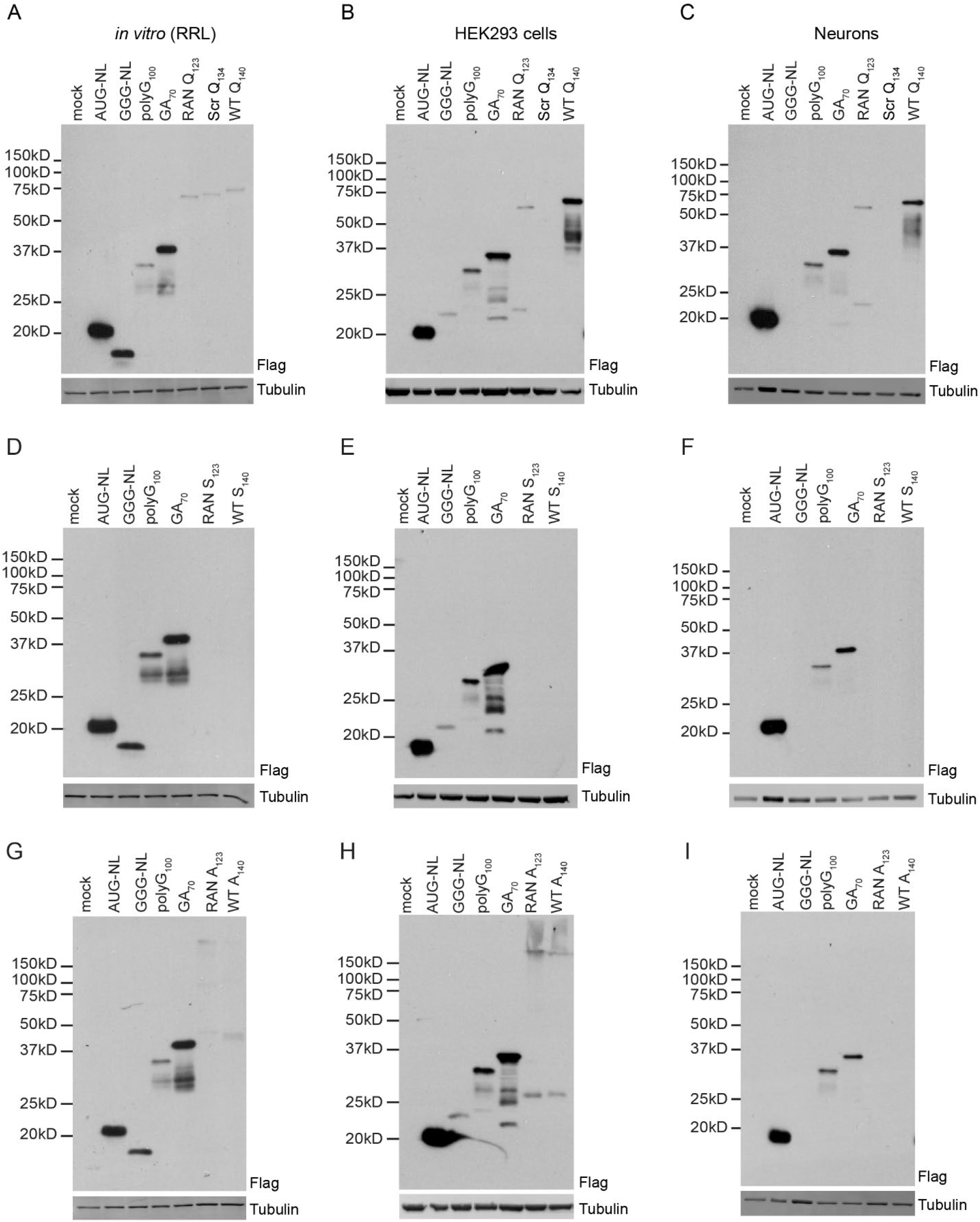
RAN reporter immunoblots derived from in vitro transcribed RNA reporters. Representative western blots from IVT RNA for the glutamine frame in RRL (A), HEK293 cells (B), or primary rodent neurons (C). Representative western blots from IVT RNA for the serine frame in RRL (D), HEK293 cells (E), or primary rodent neurons (F). Representative western blots from IVT RNA for the alanine frame in RRL (G), HEK293 cells (H), or primary rodent neurons (I). Tubulin was used as a loading control.

**Supplemental Table 1:** Start codons upstream of repeat-containing reads in RAN samples are due to sequencing artifact. Repeat containing reads from RAN samples containing upstream start codons identified using ATGFinder are listed below. Sample includes the cell type and transfected plasmid. Read ID refers to unique read coordinates and read mate. Each ATG was due to 1 or 2 base mismatches compared to reference and the position of these mismatches relative to repeat start is denoted in the position column. Two of these mismatches were near the read start and soft-clipped by the STAR aligner. Those not soft clipped were poor quality sequences where the chance of error was higher than the occurrence of the observed ATG sequence.

| Sample | Read ID | Position | Soft Clip | Phred | Sequence Error Probability | Coverage |
| --- | --- | --- | --- | --- | --- | --- |
| HEK 293<br>RAN A123 | 1290:25455:12570<br>R2 | -52 | No | 12 | 0.0631 | 4378 |
| HEK 293<br>RAN A123 | 1184:38362:29388<br>R2 | -52 | Yes | -- | -- | -- |
| HEK 293<br>RAN<br>Q123 | 1175:32900:13719<br>R2 | -31 | Yes | -- | -- | -- |
| HEK293T<br>RAN A123 | 2159:15069:22789<br>R2 | -66, -65 | No | 12, 12 | 0.0631 | 313, 315 |

